# Sinusoidal regulation denoises circadian machinery

**DOI:** 10.1101/2025.01.08.631925

**Authors:** Hotaka Kaji, Fumito Mori, Osamu Maruyama, Hiroshi Ito

## Abstract

The rhythms transmitted from the circadian clock inevitably fluctuate because of molecular noise. The level of period fluctuations, observed not in the circadian clock, but in the output system that receives the transmitted rhythm, varies according to the organism and cell type, ranging from 0.5% to 10%. However, it remains unclear how the signal transduction involved in this transmission affects the fluctuations in the oscillation period of the output system. To address this, we investigated a coupled system consisting of a circadian clock and its output. We numerically and analytically demonstrated that the rhythmic regulation through which the clock controls downstream gene expression affects the level of fluctuations in the output system. Moreover, Gibbs sampling based on the analytically obtained fluctuation formula confirmed that the sine-wave-like regulatory functions effectively minimized the fluctuation of the output system. These theoretical insights provide new perspectives on signal transduction as a denoising mechanism embodied in the circadian system.

**Author summary:** Recent single-cell observations have revealed that individual cells exhibit circadian rhythms with intrinsic variability. In particular, the period fluctuation, evaluated using the coefficient of variation (CV), was studied. In this study, we investigated how signal transduction from the central clock affects period fluctuations in the output system. We identified a key factor influencing these fluctuations: the waveform of the regulatory function by which the circadian clock governs the downstream output. We numerically demonstrated that the fluctuations vary widely depending on the regulatory function and that the sinusoidal function significantly reduced fluctuations in the output system. Furthermore, Gibbs sampling revealed that sine-like regulatory functions effectively minimized fluctuations. These findings suggest a preference for near-sinusoidal waveforms in the regulation of circadian rhythms.

## Introduction

Circadian rhythms are physiological processes that repeat approximately every 24 h. The free-running period of the circadian rhythm under constant conditions is usually reported along with its deviation from the mean oscillation period. For example, the free-running period of mice has been reported as 23.48±0.06 h [1]. Additionally, at the cellular level, the circadian rhythm of a single cellular cyanobacterium is 25.4 ±0.12 h [2] and the period of mammalian fibroblast cells is 24.38±1.12 h [3]. These deviations suggest a stochastic nature of circadian dynamics. Fluctuations in circadian rhythms originate from a self-sustained oscillator known as the circadian clock and are then transmitted to the downstream output system. Stochastic gene expression in the circadian clock machinery contributes to fluctuations in the circadian period [3–5].

Some theoretical studies have numerically and analytically addressed fluctuations in systems consisting of self-sustained oscillators [6, 7]. A formula for the period variability in general *N*-dimensional oscillatory systems has been derived [8], showing that each variable in the system can exhibit a different period variability.

The general formula for period variability was applied to a circadian clock that regulates its downstream output system [9]. This study demonstrated that the output system’s fluctuations can be smaller than those of the clock, suggesting that different organs controlled by the same oscillator can vary in their amount of fluctuation. The study considered a specific manner of signal transduction from the circadian clock by adopting sinusoidal regulation to model clock-controlled promoter activity.

However, global gene expression analyses suggest diverse ways to regulate the circadian clock [10–13]. The expression patterns of circadian-clock-controlled genes are diverse in phase, oscillation amplitude, and waveform, although these genes are controlled by a unique circadian clock. Genome-wide analysis of promoter activity using a luciferase reporter has revealed diverse waveforms (i.e., sinusoidal-like, sawtooth, and spike patterns) [14]. Variations in rhythmic patterns of gene expression can arise from differences in the sequences of clock-controlled promoters. For instance, the E-box, which is bound by the mammalian core circadian transcription factors CLOCK and BMAL1, induces daily rhythmic expression. Changes in the E-box sequence can alter the waveforms and amplitudes of the downstream output signals [15, 16]. If these variations in clock regulation are associated with fluctuations, it may be possible to control their accuracy by modifying the promoter sequences. Nonetheless, it remains unclear whether circadian clock regulation affects the level of fluctuations in downstream output.

We numerically and analytically investigated the coefficient of variation (CV, the standard deviation divided by the mean) of the oscillation period to explore the relationship between clock-controlled regulation and fluctuations in the circadian output driven by generic periodic regulation. These analyses suggested that sinusoidal regulatory functions effectively reduced downstream fluctuations.

## Results

### Regulatory function waveform influences the precision of the output rhythm

We examined the factors in the clock-regulated system that could affect the precision of the observed circadian rhythm using a coupled model of the circadian clock and its output system (Fig 1A). We employed a simple transcriptional-translational negative feedback loop (TTFL), which negatively regulates its own gene expression, as a model for the circadian clock because it is considered the basic mechanism for clocks across organisms [17]. The output system receives periodic signals from the circadian clock and converts them into specific physiological or behavioral responses. Here, we suppose reporter systems such as luciferase or fluorescent proteins as representatives of the output system with promoters regulated by a circadian clock. Additionally, we assumed that expression noise within the circadian clock system generates fluctuations in the entire circadian clock system. Based on these molecular mechanisms, we we considered the following model:

**Fig 1.**
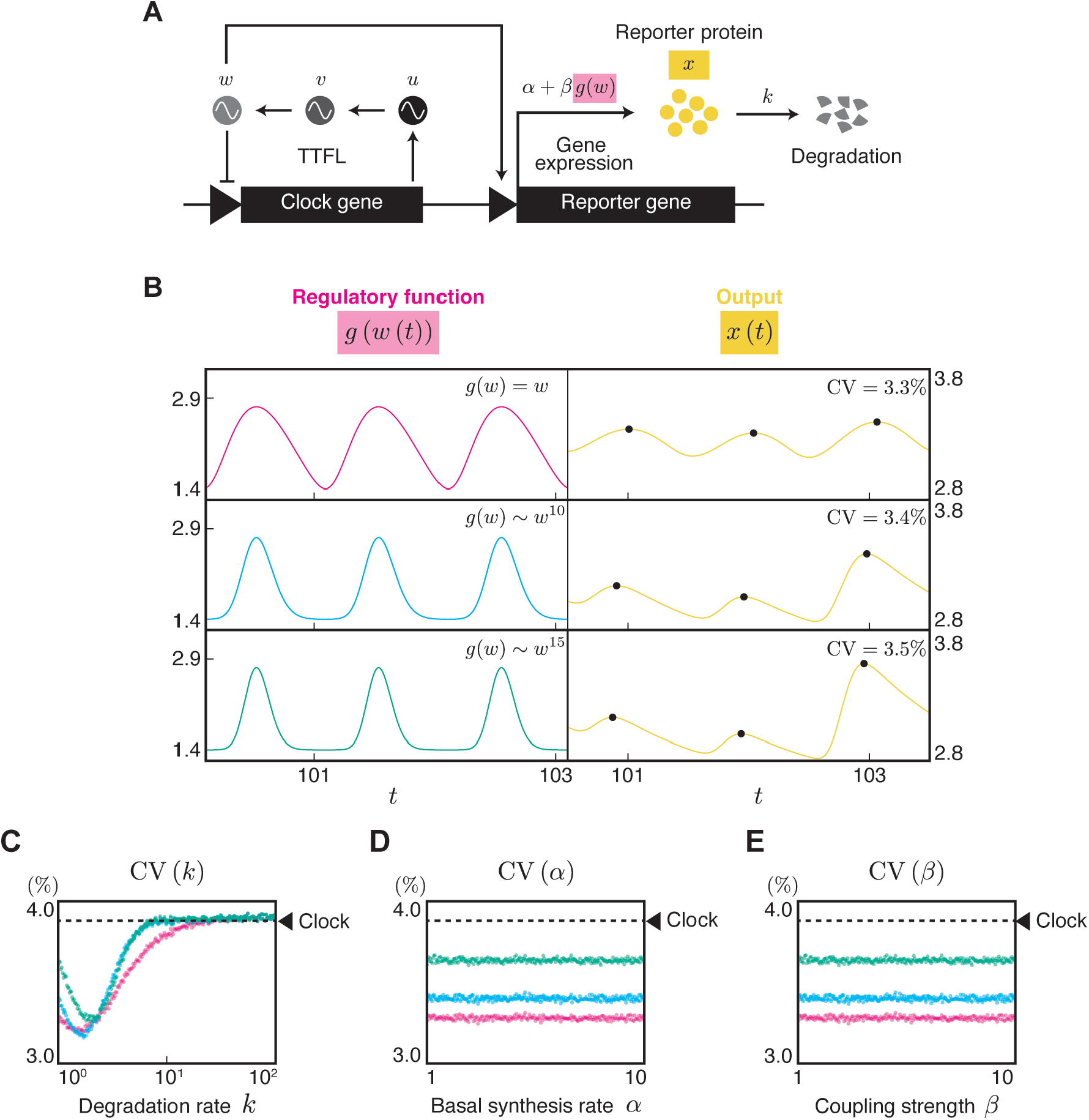
Fluctuation of oscillation period in the output system driven by TTFL. **(A)** Goodwin model representing TTFL coupled with a reporter system. The three variables in the Goodwin model, *u, v*, and *w* correspond to the amounts of mRNA, cytosolic, and nuclear clock protein, respectively. The nucleic protein *w* as a circadian clock drives the promoter activity of the reporter gene via the regulatory function *g*(*w*). Thus, the reporter protein *x* is rhythmically expressed. The reporter protein degrades at a rate of *k*. Gene expression noise causes fluctuations in *u*, resulting in fluctuations throughout the entire system. (**B**) Different regulatory functions lead to different fluctuations in the output system. The left column shows the time course of *g*(*w*(*t*)) where *ϵ* = 0. The three different regulatory functions represent *g*(*w*) = *w* (red), *g*(*w*) = 4.558×10^−5^*w*^10^ + 1.555 (blue), and *g*(*w*) = 2.783×10^−7^*w*^15^ + 1.559 (green). The right column shows *x*(*t*) obtained through numerical simulations for Eq (1) using corresponding *g*(*w*). The black dots indicate the peaks of *x*(*t*). The parameters in the simulation were set to *k*_*u*_ = *k*_*v*_ = *k*_*w*_ = 0.1, *α* = *β* = 1, *k* = 1, *ϵ* = 10^−4^, *D* = 1.0, and *τ* = 39.7. (**C**) The dependency of the degradation rate *k* on the CV of the oscillation period. *α* = *β* = 1. (**D**) The dependency of the basal synthesis rate *α* on the CV of the oscillation period. *β* = 1, *k* = 1. (**E**) The dependency of the coupling strength *β* on the CV of the oscillation period. *α* = 1, *k* = 1. The colors correspond to the functions in Fig B. The values of the other parameters are the same as in Fig B. The dashed line indicates the fluctuation in the period of the nucleic clock protein *w*.

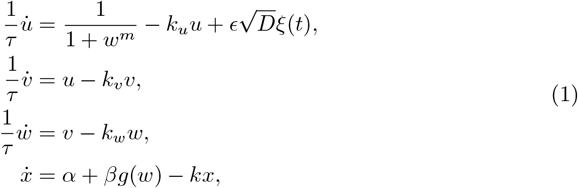

where *u* represents the amount of clock gene mRNA, *v* and *w* represent the amount of clock proteins in the cytoplasm and nucleus, respectively, and *x* represents the amount of reporter proteins as output system. The coefficient *ϵ* (⪡1) is a small parameter, 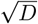 is the noise intensity of gene expression of the clock gene, and *ξ*(*t*) is an independent Gaussian noise satisfying *E*[*ξ*(*t*)] = 0, *E*[*ξ*(*t*)*ξ*(*t*′)] = *δ*(*t−t*′), where *E*[·] represents the expectation and *δ*(*t*) is the Dirac delta function. *m* denotes the Hill coefficient and *k*_*u*_, *k*_*v*_, *k*_*w*_, *k* are the degradation rates of each molecule. The first three equations are the Goodwin model with three variables [18, 19] representing TTFL, where the production rate 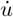 of the clock gene is negatively regulated by the clock protein *w*. The TTFL can show a self-sustained oscillation. We introduce the scale parameter for time *τ*, which is the original period of the limit cycle of the Goodwin model in the absence of noise (*ϵ* = 0), such that the mean oscillation period of Eq (1) is unity, corresponding to a day. A clock protein rhythmically controls the synthesis of reporter proteins at a rate *α* + *βg*(*w*). Simultaneously, the reporter protein is degraded at a rate of *k*. When *ϵ* ≠ 0, *w* oscillates with fluctuations. Consequently, the synthesis controlled by *w* also fluctuates, leading to oscillations with varying periods in the output *x*. Here, we assume that *α* represents the basal synthesis rate of the reporter protein and *β* represents the coupling strength between the clock and downstream gene regulation. Function *g*(*w*) describes the regulation by the clock protein *w* on reporter gene expression.

We first investigated the effect of the waveform of the regulatory function *g*(*w*) on the fluctuations in the output (Fig 1B). Three types of polynomial functions were assigned to *g*(*w*): first-degree, tenth-degree, and fifteenth-degree polynomials. A higher degree makes the waveform of *g*(*w*) sharper. We performed numerical simulations of Eq (1) under different regulatory functions *g*(*w*) and then quantified the CV of the peak-to-peak period (Numerical simulation, §4.1 in the Supplementary text). As shown in Fig 1B, different rhythmic regulations *g*(*w*) lead to varying levels of fluctuations in the output system *x*(*t*), even though the output systems are driven by the same clock. The regulatory function with a sharper waveform showed a higher CV and larger amplitude variation.

We also investigated whether factors other than *g*(*w*) influenced CV in the reporter system. The degradation rate of the reporter protein *k* affected the amount of fluctuation (Fig 1C). Within the range of *k*, the fluctuations were smaller than those inherent to the original circadian clock. This observation is consistent with that of Kaji et al. [9]. In contrast, the basal synthesis rate *α* and coupling strength *β* did not influence the fluctuations in the output system (Figs 1D and 1E). The dependence of *k* and the independence of *α* and *β* on the output fluctuations were consistent regardless of the type of *g*(*w*).

In §2 of the Supplementary text, we examined the case in which *u* or *v* regulates the output system instead of *w*. As Fig S1B shows, the CV of the output system depended on the choice of *g*(*u*), *g*(*v*), or *g*(*w*). Moreover, the dependence of *k* on CV as well as the independence of *α* and *β* remained consistent regardless of the regulatory function employed.

### Sinusoidal regulation for the output system maintained a precise circadian rhythm

Recent findings on the molecular mechanism of the circadian clock indicate that the circadian clock system is a more complex network than a single transcriptional-translational loop, that is, multiple gene regulatory loops [20, 21] or a combination of biochemical oscillators and TTFL [22, 23]. The applicability of results obtained solely from the Goodwin model is limited to real circadian clocks. Therefore, we adopted a more general model, the phase oscillator, as the circadian clock model, which controls the reporter system through regulatory function *f* (Fig 2A):

**Fig 2.**
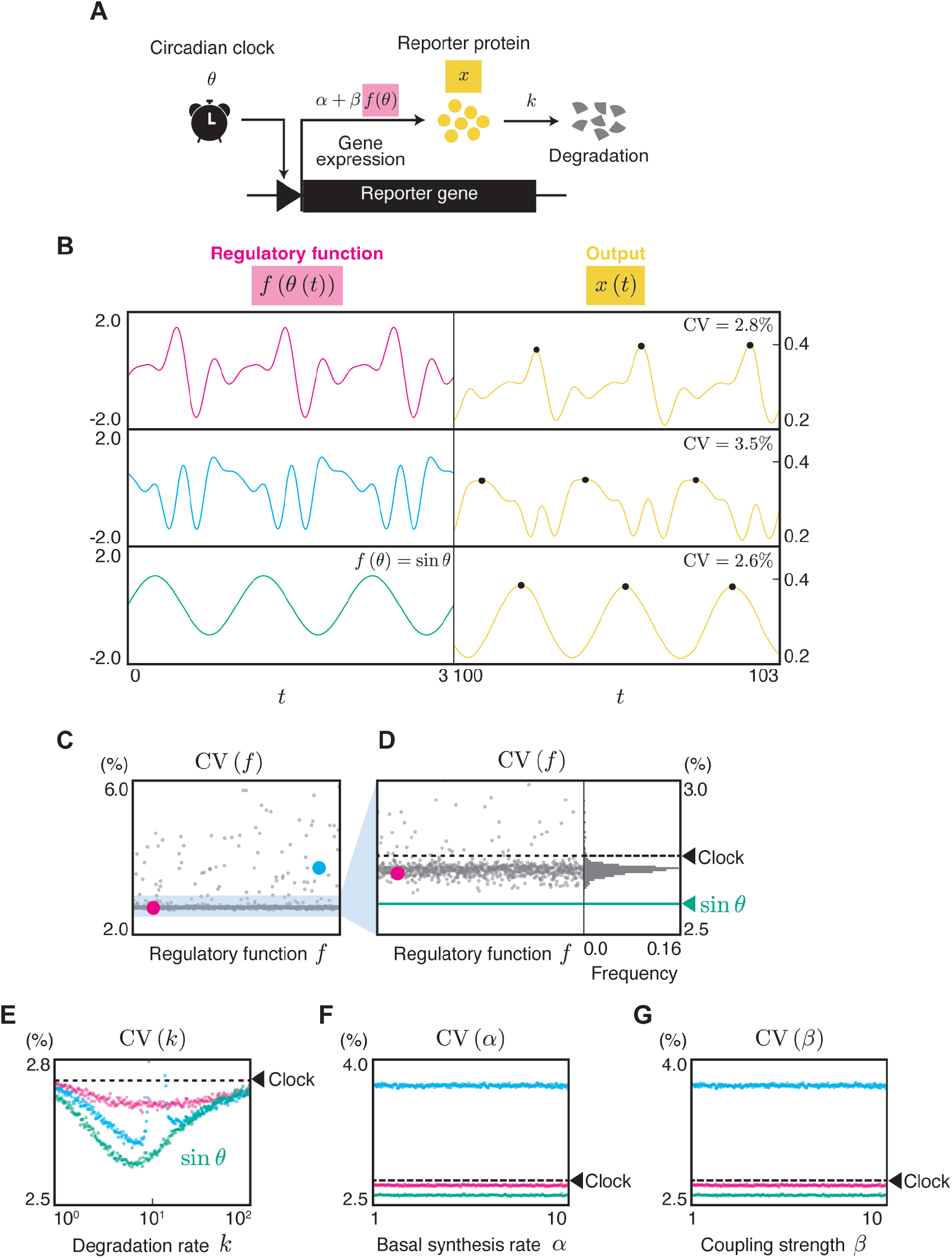
Fluctuation of oscillation period in the output system driven by a phase oscillator. (**A**) Reporter system driven by a phase oscillator as a general model for the circadian system. The phase oscillator rhythmically controls the reporter gene through a clock-controlled promoter at a rate *α* + *βf* (*θ*). The expressed protein is degraded at a rate of *k*. (**B**) Different regulatory functions lead to different fluctuations in the output system. The left column shows the time course of *f* (*θ*(*t*)) where *ϵ* = 0. The right column shows *x*(*t*) obtained through numerical simulations for Eq (2) using corresponding *f* (*θ*). The black dots indicate the peaks of *x*(*t*). The parameters in the simulation were set to *α* = 3, *β* = 1, *k* = 10, *ω* = 2*π, ϵ* = 0.1, *D* = 3.0. The Fourier coefficients of red and blue waveforms are shown in §6 of the Supplementary text. (**C, D**) CV of the oscillation period of *x* for randomly chosen *f* obtained from numerical simulation. Fig D is the magnified plot for Fig C with a histogram. The CV for the sinusoidal regulation *f* (*θ*) = sin *θ*, represented by the green line, is 2.6%. (**E**) The dependency of the degradation rate *k* on the CV of the oscillation period. The colors correspond to the functions in Fig B. (**F, G**) The dependency of *α* and *β* on the CV of the oscillation period, respectively. We assigned values for the fixed parameters as follows: *α* = 3, *β* = 1, *k* = 10 for Figs C and D; *α* = 1, *β* = 0.4 for Fig E; *β* = 0.4, *k* = 10 for Fig F; *α* = 23, *k* = 10 for Fig G; the other parameters are commonly set to *ω* = 2*π, ϵ* = 0.1, *D* = 3.0. In Figs D, E, F, and G, the dashed line represents the CV of the clock, expressed as 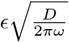 [9].

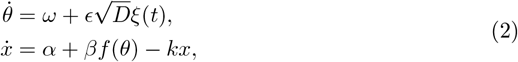

where *θ*(*t*) is the phase of the circadian clock (modulo 2*π*) and *ω* is the angular frequency of the clock. The function *f* (*θ*) describes the regulation of reporter gene expression by clock phase *θ*. The reporter protein is synthesized at a rate of *α* + *βf* (*θ*). The other parameters correspond to those in Eq (1). Note that the protein synthesis rate *α* + *βf* (*θ*) ≥ 0 must hold. Eq (2) has a limit cycle solution with a period *τ* = 2*π/ω* when *ϵ* = 0. When *ϵ* ≠ 0, the trajectory of Eq (2) fluctuates around the limit cycle owing to noise.

To facilitate analytical and numerical calculations for this model, we expanded

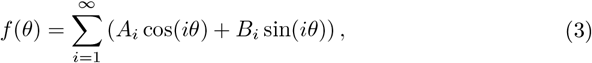

where *i* denotes a harmonic number. The Fourier coefficients were normalized to 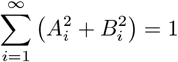. *A*_1_ cos *θ* and *B*_1_ sin *θ* reflect the primary oscillatory behavior corresponding to 24-h cycle; hence, *A*_1_ or *B*_1_ should be nonzero. Higher-order terms (*i >* 1) in the Fourier expansion represent additional harmonics that contribute to more complex waveforms, such as sharper peaks or broader troughs in the oscillations. These higher harmonics allowed the model to capture deviations from a simple sine curve.

First, we numerically investigated whether the CV in the output system of Eq (2) depends on the factors *f* (Numerical simulation, §4.1 in the Supplementary text). We measured the CV of the peak-to-peak intervals of *x* in the model with three different regulatory functions (Fig 2B). These three regulatory functions resulted in different CVs. This observation is consistent with that shown in Fig 1.

For further investigation, we observed a CV with randomly generated 1,000 regulatory functions *f* that were expressed as a 5th order Fourier series:

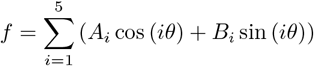,where *A*_*i*_, *B*_*i*_ ∈ [−1, 1]. The different regulatory functions resulted in a wide range of CV values, and most CV values are lower than the CV of the circadian clock (Figs 2C and 2D). As shown in Fig 2D, none of the randomly generated functions exhibited smaller fluctuations than *f* (*θ*) = sin *θ*. This implies that sinusoidal regulation significantly reduces fluctuations.

In addition, we investigated the CV under varying values of *α, β*, and *k*, focusing on the three selected functions (red, blue, and green in Fig 2B). The degradation rate *k* influenced the fluctuations in the output system (Fig 2E) regardless of the regulatory function *f*, yet the extent and direction of fluctuation enhancement or reduction were dependent on the specific regulatory function. In contrast, parameters *α* and *β* showed no significant effect on CV, regardless of *f* (Figs 2F and 2G). These findings align with the Goodwin model (Figs 1C, 1D, and 1E).

Random sampling of *f* is not sufficient to properly evaluate the predominant denoising effect of sinusoidal regulation, owing to the high-dimensional parameter space. Thus, we performed Gibbs sampling to effectively seek functions with a high denoising ability (Fig 3A) (see Gibbs sampling in the Methods section). The distribution of 20,000 samples from Gibbs sampling successfully shifted towards a lower CV compared with random sampling. The mean value of the sampled CV shifted from 3.0% to 2.7% and the minimum shifted from 2.64% to 2.56%. Despite the shift in the distribution of CV, most regulatory functions sampled through Gibbs sampling still exhibited a higher CV than sinusoidal regulation, supporting the strong noise reduction ability of sinusoidal regulation. Only 0.3% of the sampled functions exhibited a lower CV than the sinusoidal function.

**Fig 3.**
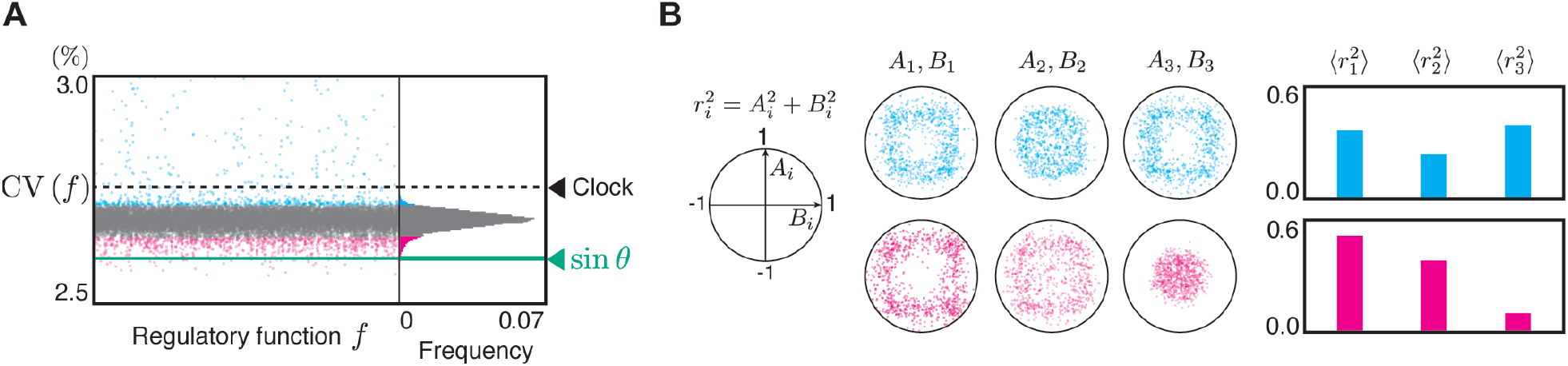
Gibbs sampling for regulatory functions to show lower CV obtained by numerical simulation. (**A**) The samples obtained by Gibbs sampling, with only those having CVs not exceeding 3.0% being shown. The corresponding histogram is displayed on the right. The number of Fourier harmonics for regulatory functions was set to 3. *ω* = 2*π, ϵ* = 0.1, *D* = 3.0, *α* = 3, *β* = 1, *k* = 10. The dashed line represents the CV of the clock, expressed as 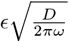 [9]. The CV for the sinusoidal regulation *f* (*θ*) = sin *θ*, represented by the green line, is 2.6%. (**B**) Distribution of Fourier coefficients for the regulatory function *f* (*θ*), corresponding to the fluctuations represented by the red and blue dots in Fig A. Red and blue indicate the top and worst 5% of all samples from Gibbs sampling. Note that each unit circle, defined by 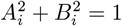, is an upper boundary because of the normalization of Fourier coefficients. For further details on Gibbs sampling see the Methods section

The Fourier coefficients of the functions obtained through Gibbs sampling show distinct distributions depending on the CV value (Fig 3B). Furthermore, we examined the average power of the Fourier coefficients for different harmonic numbers *i*, defined as 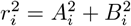,across the sampled functions denoted by 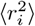.For functions exhibiting higher CV, there was little difference in the values of 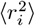 across the different harmonic numbers *i*. In contrast, functions exhibiting a lower CV show larger values of 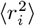 for smaller harmonic numbers *i*. This suggests that the sampled functions exhibiting lower CV were closer to sinusoidal shapes in Fourier space. In §5 of the Supplementary text, we confirm that a sine-like function achieves the lowest output fluctuation by performing evolutionary optimization in the fifth-order Fourier coefficient space.

### Analytical calculations supporting the results obtained from numerical simulations

To mathematically ensure the dependence of *f* and *k* and the independence of *α* and *β* on CV shown in the previous section, we analytically calculated the fluctuation in the period indicated by *x*(*t*) in Eq (2) based on the theory proposed by Mori & Mikhailov [8]. This theory provides the analytical way to calculate CV of the oscillation period defined by checkpoints which correspond to the biological concept of “gating” in chronobiology. The CV of a general *N*-dimensional oscillatory system can be expressed as follows:

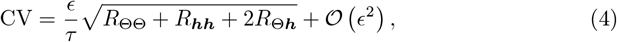

where *R*_ΘΘ_ is the variance of the system-level phase, *R*_***hh***_ is the autocorrelation of the amplitude deviation, and *R*_Θ***h***_ is the cross-correlation between the system-level phase shift and amplitude deviation. *R*_ΘΘ_ and *R*_***hh***_ are always nonnegative, but *R*_Θ***h***_ can become negative. When *ϵ* is sufficiently small, CV can be expressed using only the three aforementioned components.

For the analytical derivation of the CV of the output system in Eq (2) using the theory [8], an explicit expression of the limit cycle for Eq (2) is necessary. Because we can derive the limit cycle solution analytically, we can compute the three components of Eq (4) for Eq (2) analytically:

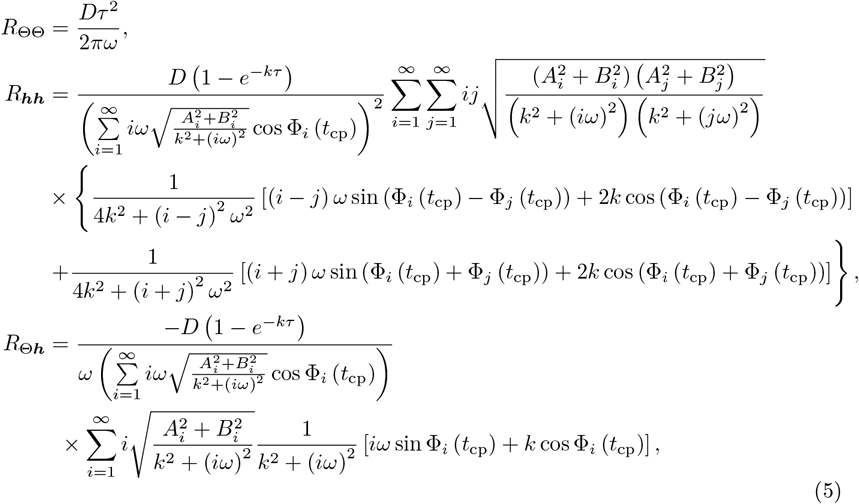

Where 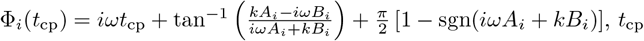 is a time parameter related to checkpoint (see §4.2 in the Supplementary text), and sgn is the sign function.

We find that all three fluctuation factors *R*_ΘΘ_, *R*_***hh***_, and *R*_Θ***h***_ are independent of *α* and *β*, while *R*_***hh***_ and *R*_Θ***h***_ are dependent on *k* and (*A*_*i*_, *B*_*i*_), i.e., *f*. In other words, our analysis, despite not employing peak-to-peak intervals, supports the *k* and *f* dependencies and the *α* and *β* independencies of the CV, which were observed in the numerical results based on the peak-to-peak intervals.

Fig 4 confirms the agreement between the numerical simulations and analytical calculations. In particular, the agreement strengthened under smaller noise. Here, we employed the midpoint of the oscillation range instead of the peaks to define the period (see §4 in the Supplementary text for details).

**Fig 4.**
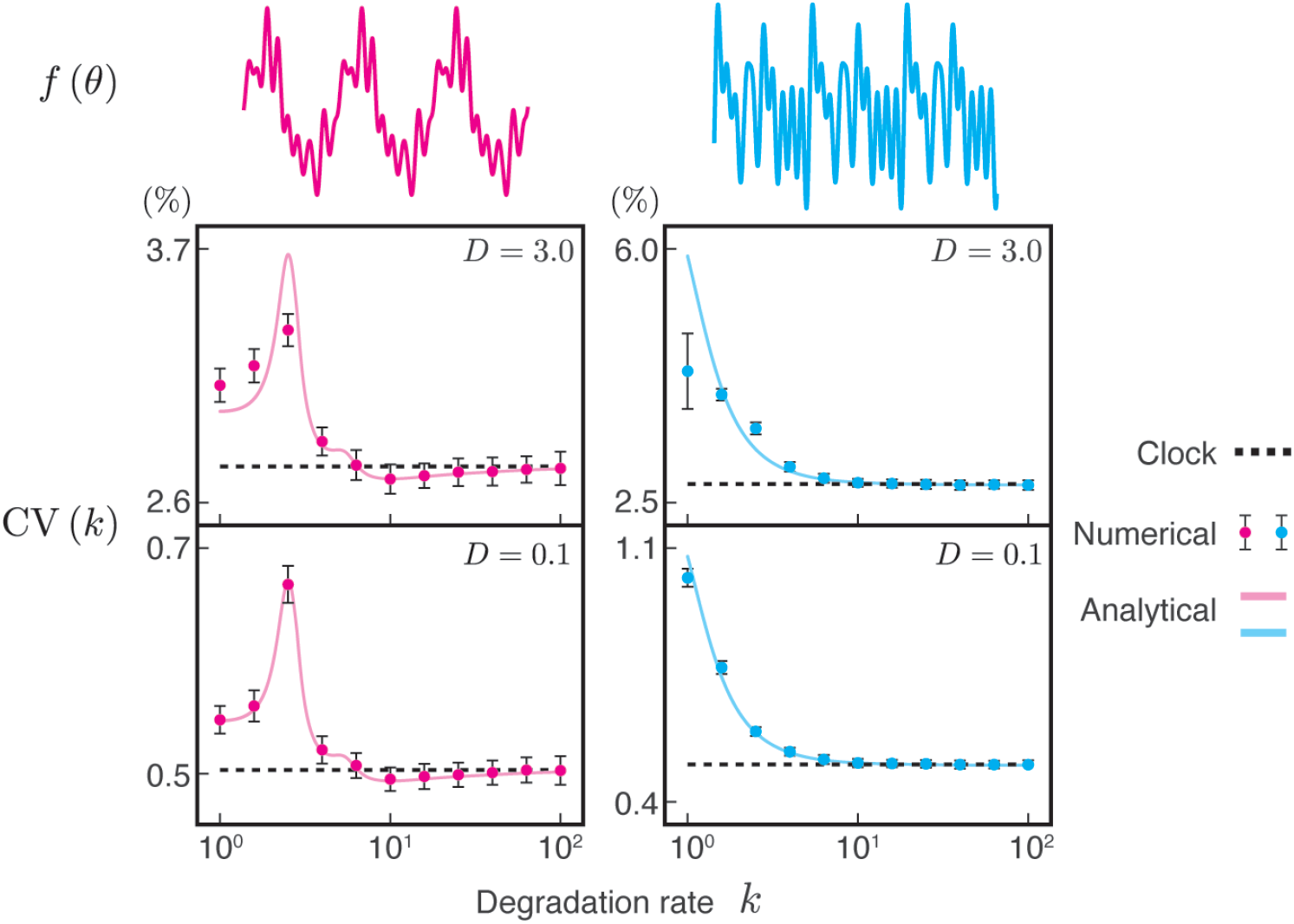
Numerical and analytical fluctuations in the oscillation period. The CV of the oscillation period of the reporter protein under two different clock regulations (red and blue). The Fourier coefficient of each *f* (*θ*) is shown in §6 in the Supplementary text. The period was measured based on the threshold (see §4 in Supplementary text). *ω* = 2*π, ϵ* = 0.1, *α* = 4, and *β* = 1.

Using the formula Eq (4) together with Eq (5) for the output system of Eq (2), we performed Gibbs sampling to explore the regulatory function *f* which further reduces fluctuations without requiring numerical simulations (see Gibbs sampling in the Methods section). This analytical approach has the advantage of a significantly lower computational cost than numerical simulations. Consequently, we obtained many more functions that exhibited a lower CV than the numerical approach (Fig 5A). Most of the sampled functions (93.3%) yielded a higher CV than the sine function, whereas some functions belonging to a small fraction resulted in a lower CV. The Fourier coefficients of the functions derived through Gibbs sampling show distinct distributions depending on the CV value (Fig 5B). Functions exhibiting CVs larger than the clock showed little difference in 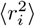 across different harmonic numbers.

**Fig 5.**
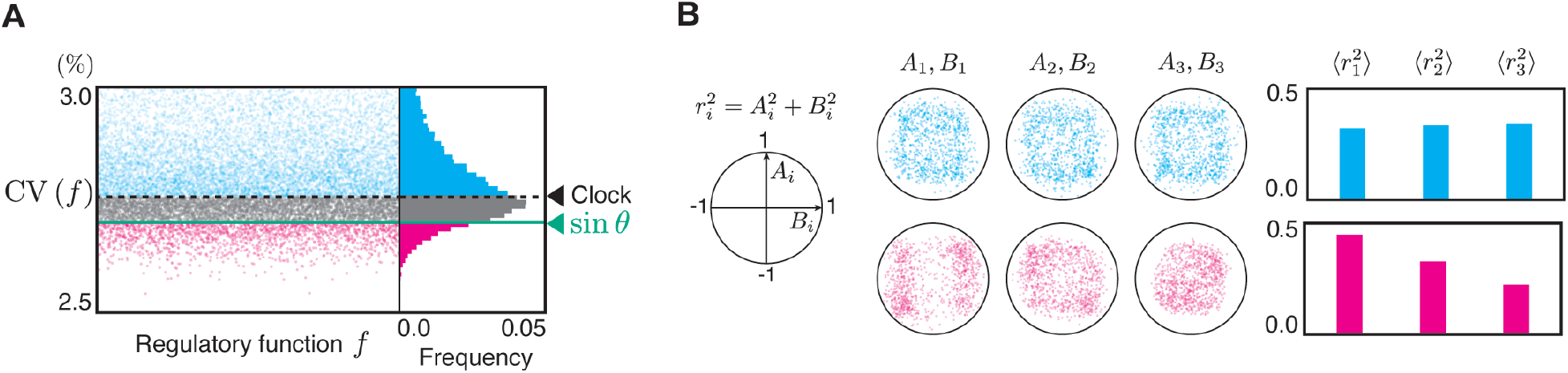
Gibbs sampling for the regulatory functions to show low CV obtained by the analytical formula of CV. (**A**) We obtained 10^7^ regulatory functions through Gibbs sampling. To prevent the display from becoming overcrowded with too many points, we reduced the number of points by downsampling and displayed only 0.2% of the sampled functions. The CV for the sinusoidal regulation, represented by the green line, is 2.7% (see §4 in the Supplementary text). The red and blue dots represent functions exhibiting CVs higher than the CV of the clock and lower than those of the sine, respectively. The right panel shows the histogram corresponding to the left panel. The analytically obtained CV is calculated based on periods defined by a threshold (see §4 in the Supplementary text). The dashed line represents the CV of the clock, expressed as 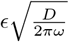 [9]. The parameters are *ω* = 2*π, ϵ* = 0.1, *D* = 3.0, and *k* = 10. (**B**) Distribution of Fourier coefficients for the regulatory function *f* (*θ*), corresponding to the fluctuations represented by the red and blue dots in Fig A. Note that each unit circle, defined by 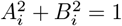 is an upper boundary because of the normalization of Fourier coefficients. A set of 1000 functions was randomly chosen, each with higher CVs than the clock and lower than the sine.

Meanwhile, functions with CVs smaller than *f* (*θ*) = sin *θ* showed larger values of 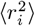 for smaller harmonic numbers, which is consistent with the results of the Gibbs sampling of numerically obtained CVs (Fig 3B). These findings confirmed the high denoising effect of the sinusoidal regulatory function as demonstrated by the numerical approach. Indeed, the functions exhibiting a lower CV appeared closely sinusoidal in §3 of the Supplementary text.

## Discussion

In this study, we focused on the amount of fluctuations transmitted from the circadian clock to the output system (Figs 1 and 2). These fluctuations were found to depend on the gene expression regulatory functions (*g* in Fig 1 and *f* in Fig 2) and the degradation rate, but not on the basal synthesis rate or coupling strength. Such dependence and independence were found not only in the Goodwin model Eq (1) but also in the phase model Eq (2), reinforcing the generality of these characteristics. Therefore, we expect these characteristics to apply to real systems.

In our numerical simulations, we limited the harmonic number of the Fourier series of the regulatory function to finite values. Here, we validated whether this limitation was appropriate for the regulatory function form that exhibited a lower CV. As shown in Figs 3B and 5B, functions exhibiting lower CV tend to have smaller values of Fourier coefficients *A*_*i*_ and *B*_*i*_ as the harmonic number *i* increases. Similarly, this tendency was observed in the optimal functions obtained using an evolutionary algorithm to minimize the CV, where functions with higher-frequency components were employed (Figs S5 and S6). These results indicated that the high-frequency components of the regulatory function did not contribute significantly to reducing CV. Therefore, our analysis, which employed functions with a finite number of Fourier harmonics, is valid for identifying waveforms with lower CV.

Our numerical simulation revealed the potential of sine functions to enhance the precision of the output systems. In reality, the circadian network may play a significant role in reducing fluctuations because the observed rhythms often closely resemble a sine curve [2, 3]. As shown in Figs 1B, 2B, S1B, and S6B, the waveforms of the output system *x*(*t*) tended to resemble those of its corresponding regulatory functions. This suggests that the observed circadian rhythms resemble the waveforms of their regulatory functions. The resemblance of the observed circadian rhythms to a sine curve, combined with the assumption that the output waveforms reflect their regulatory functions, indicates that the real regulatory functions of circadian rhythms resemble sinusoidal patterns. Our findings demonstrated that sinusoidal regulation effectively reduced fluctuations and enhanced the precision of the output system, suggesting a predominance of the actual observed sine-like waveforms of circadian rhythms in terms of controlling fluctuations. In contrast, the action potentials of neurons [24] and cardiac muscle cells [25] exhibited sharp waveforms. Neurons and cardiac muscle cells may not require highly precise rhythms compared with the circadian machinery.

Experimentally altering the waveforms of circadian rhythms provides a means to verify whether the waveforms of regulatory functions govern fluctuations in actual circadian systems because the waveforms of observed circadian rhythms and the corresponding regulatory functions are expected to be similar. At a physiological level, circadian oscillations pass through various promoters originating from a single circadian clock [26]. For instance, the waveforms of circadian rhythms can be manipulated by modifying circadian regulatory promoter sequences [14] or introducing time delays into feedback loops [27]. Additionally, altering signal transduction pathways may alter circadian rhythm waveforms. For example, bypassing another downstream pathway can result in bimodal oscillations [28]. Furthermore, changes in waveforms owing to temperature variations have been reported, with observations in cyanobacteria showing that the circadian rhythm waveform approaches a sine wave under low-temperature conditions [29]. It is possible to confirm whether changes in the waveform of the regulatory functions by these factors affect the level of fluctuations in the output system.

Experimental chronobiology researchers have focused on the average circadian rhythm period. Screening for mutants with irregular average periods led to the identification of several clock genes. The molecular networks consisting of these clock genes are the core circadian clocks [17]. In contrast, this study focused on the variance in the periods and waveforms of circadian rhythms, which has attracted little attention in chronobiology. Screening based on period variance and waveforms can lead to the discovery of unknown clock genes. Furthermore, the effects of mutations in clock genes on the period variance and waveforms remain unexplored, except in a few studies [23, 30]. The present study, which elucidated the relationship between period variance and waveforms, not only highlights novel functions of waveforms in self-sustained oscillators (e.g., [31–33]) but also marks a milestone in expanding the scope of chronobiology.

## 1 Methods

### Numerical simulation

We numerically solved Eqs (1) and (2) using the Euler-Maruyama method. The initial conditions of Eq (2) were set on the limit cycle (§1 in the Supplementary text) as 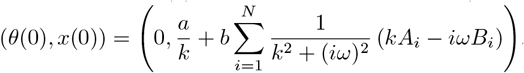. For the Goodwin model, Eq (1), the initial conditions were set to (*u*(0), *v*(0), *w*(0), *x*(*t*)) = (0, 0, 0, 0). The period was measured from *t* = 100. For both the simulations, the time step was Δ*t* = 1.0 × 10^−3^.

### Gibbs sampling

Gibbs sampling was performed to investigate the distribution of regulatory function *f*, which showed smaller fluctuations. The Fourier coefficients of *f*, (*A*_1_, *B*_1_, *A*_2_, *B*_2_, *A*_3_, *B*_3_), are denoted as **x** = (x_1_, x_2_, x_3_, x_4_, x_5_, x_6_). Initially, a random value from a uniform distribution in the range [− 1, 1] was assigned to x_*i*_ (*i* = 1, 2, …, 6). Then, for *i* = 1, 2, …, 6, the value of x_*i*_ is updated according to the conditional probability distribution

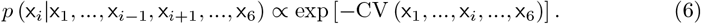

To numerically evaluate the conditional probability distribution *p*, the value of Eq (6) was calculated for 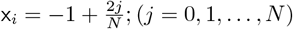.This procedure was repeated until *n*_max_ samples were obtained. The first 1000 samples were discarded during the burn-in period. *n*_max_ = 21000 and *N* = 50 for Fig 3. *n*_max_ = 10^7^ and *N* = 1000 for Fig 5.

## Data Availability Statement

All programming codes are publicly available on GitHub at https://github.com/hito1979/outputnoise.

## Acknowledgments

We sincerely thank Hideyuki Takagi (Kyushu University) for engaging discussions and valuable advice on Differential Evolution.

## Author Contributions

**Conceptualization:** Hotaka Kaji, Mori Fumito, Hiroshi Ito.

**Formal analysis:** Hotaka Kaji, Mori Fumito, Osamu Maruyama, Hiroshi Ito.

**Writing – original draft:** Hotaka Kaji, Mori Fumito, Hiroshi Ito.

**Writing – review & editing:** Hotaka Kaji, Mori Fumito, Hiroshi Ito.

