## Supplementary Text for "Sinusoidal regulation denoises circadian machinery"

#### Contents

**1 Derivation of Eq (5)** **1**

**2 Regulation by the different components in the Goodwin model Eq (1)** **3**

**3 Examples of regulatory functions obtained by Gibbs sampling** **4**

**4 Methods to obtain CV of period** **5**
4.1 CV obtained from numerical simulations . . . . . . . . . . . . . . . . . . . . . . . . . 5
4.2 CV obtained from Eq (5) . . . . . . . . . . . . . . . . . . . . . . . . . . . . . . . . . . 6

**5 Optimization of the regulatory function via an evolutionary algorithm** **7**

**6 Supplementary table** **10**

### 1 Derivation of Eq (5)

Based on the theory formulated by Mori & Mikhailov [1], we analytically derived the fluctuation in the oscillation period of the model,

$$\begin{aligned} \dot{\theta} &= \omega + \epsilon\sqrt{D}\xi(t), \\ \dot{x} &= \alpha + \beta f(\theta) - kx. \end{aligned} \tag{S1}$$

This theory provides a method for analytically calculating the fluctuations in the oscillation period of a general $N$-dimensional oscillatory system in the presence of noise described as:

$$\frac{d\boldsymbol{x}}{dt} = \boldsymbol{F}[\boldsymbol{x}(t)] + \epsilon\boldsymbol{G}[\boldsymbol{x}(t)]\boldsymbol{\xi}(t),$$

where $\boldsymbol{x}(t)$ is a vector with $N$ elements, $\boldsymbol{F}$ represents the $N$-dimensional oscillatory system that generates a limit-cycle solution with the period $\tau$ under noiseless conditions, $\epsilon \ll 1$ is a small parameter, $\boldsymbol{G}[\boldsymbol{x}(t)]$ is an $N \times N$ diagonal matrix, and $\boldsymbol{\xi}(t)$ is a vector of additive noise, with $E[\xi_i] = 0$, $E[\xi_i(t_1)\xi_j(t_2)] = \delta_{ij}\delta(t_1 - t_2)$; $\delta_{ij}$ is the Kronecker delta. For the model in Eq (S1), $\boldsymbol{x}$, $\boldsymbol{F}$, $\boldsymbol{G}$, and $\boldsymbol{\xi}$ can be written as

$$\begin{aligned} \boldsymbol{x}(t) &= \begin{bmatrix} \theta(t) \\ x(t) \end{bmatrix}, \\ \boldsymbol{F}[\boldsymbol{x}(t)] &= \begin{bmatrix} F_\theta(\theta, x) \\ F_x(\theta, x) \end{bmatrix} = \begin{bmatrix} \omega \\ \alpha + \beta f(\theta) - kx \end{bmatrix}, \\ \boldsymbol{G}[\boldsymbol{x}(t)] &= \begin{bmatrix} \sqrt{D} & 0 \\ 0 & 0 \end{bmatrix}, \\ \boldsymbol{\xi}(t) &= \begin{bmatrix} \xi(t) \\ 0 \end{bmatrix}. \end{aligned} \tag{S2}$$

$f(\theta)$ can be expanded as shown in Eq (3). The solution to Eq (S1) in the absence of noise, that is, $\epsilon = 0$, is

$$\theta(t) = \omega t + \theta(0) \mod 2\pi,$$
$$x(t) = \frac{\alpha}{k} + \beta \sum_{i=1}^{\infty} \frac{1}{k^2 + (i\omega)^2} \left[ (i\omega A_i + kB_i) \sin\left[ i\left(\omega t + \theta(0)\right)\right] + (kA_i - i\omega B_i)\cos\left[ i\left(\omega t + \theta(0)\right)\right]\right]$$
$$+ \left\{ x(0) - \frac{\alpha}{k} - \beta \sum_{i=1}^{\infty} \frac{1}{k^2 + (i\omega)^2} \left[ (i\omega A_i + kB_i) \sin\left(i\theta(0)\right) + (kA_i - i\omega B_i)\cos\left(i\theta(0)\right)\right]\right\} e^{-kt}.$$

Then, if we choose a point on the limit cycle, $(\theta(0), x(0)) = \left(0, \frac{\alpha}{k} + \beta \sum\limits_{i=1}^{\infty} \frac{1}{k^2+(i\omega)^2}\left(kA_i - i\omega B_i\right)\right)$ as the initial condition, the limit cycle solution, satisfying $\boldsymbol{p}(t) = \boldsymbol{p}(t+\tau)$, is described as

$$\boldsymbol{p}(t) = \begin{bmatrix} p_1(t) \\ p_2(t) \end{bmatrix} = \begin{bmatrix} \omega t \mod 2\pi \\ \frac{\alpha}{k} + \beta \sum\limits_{i=1}^{\infty} \sqrt{\frac{A_i^2+B_i^2}{k^2+(i\omega)^2}} \sin \Phi_i\left(t\right) \end{bmatrix} \tag{S3}$$

where $\Phi_i(t) = i\omega t + \tan^{-1}\left(\frac{kA_i - i\omega B_i}{i\omega A_i + kB_i}\right) + \frac{\pi}{2}\left[1 - \mathrm{sgn}(i\omega A_i + kB_i)\right]$, and sgn denotes the sign function.

Assuming that $\boldsymbol{x}(t)$ is near the limit cycle trajectory, $\boldsymbol{x}(t)$ can be written as:

$$\boldsymbol{x}(t) = \boldsymbol{p}(t) + \epsilon \boldsymbol{z}(t) + \boldsymbol{O}(\epsilon^2),$$

where $||\boldsymbol{z}|| \ll \epsilon^{-1}$, In this case, $\boldsymbol{z}(t)$ follows the linearized equation:

$$\frac{d\boldsymbol{z}}{dt} = \boldsymbol{\Gamma}(t)\boldsymbol{z}(t) + \boldsymbol{G}[\boldsymbol{p}(t)]\boldsymbol{\xi}(t).$$

For Eq (S1), the Jacobian matrix $\boldsymbol{\Gamma}(t)$ is:

$$\boldsymbol{\Gamma}(t) = \begin{bmatrix} 0 & 0 \\ \beta \sum\limits_{i=1}^{\infty} i\left(-A_i \sin\left(i\omega t\right) + B_i \cos\left(i\omega t\right)\right) & -k \end{bmatrix}.$$

Next, we considered an unperturbed system:

$$\frac{d\boldsymbol{z}}{dt} = \boldsymbol{\Gamma}(t)\boldsymbol{z}(t) = \begin{bmatrix} 0 & 0 \\ \beta \sum\limits_{i=1}^{\infty} i\left(-A_i \sin\left(i\omega t\right) + B_i \cos\left(i\omega t\right)\right) & -k \end{bmatrix} \begin{bmatrix} z_1(t) \\ z_2(t) \end{bmatrix}. \tag{S4}$$

Eq (S4) can be solved using the initial value $\boldsymbol{z}(0)$ as follows:

$$\boldsymbol{z}(t) = \begin{bmatrix} 1 & 0 \\ \beta \sum\limits_{i=1}^{\infty} \frac{i}{k^2+(i\omega)^2}\left[\sqrt{\left(A_i^2+B_i^2\right)\left(k^2+(i\omega)^2\right)}\cos\Phi_i(t) - \left(i\omega A_i + kB_i\right)e^{-kt}\right] & e^{-kt} \end{bmatrix} \begin{bmatrix} z_1(0) \\ z_2(0) \end{bmatrix}.$$

Moreover, $\boldsymbol{z}(t)$ can be rewritten as:

$$\boldsymbol{z}(t) = \boldsymbol{U}(t)\boldsymbol{U}(0)^{-1}\boldsymbol{z}(0),$$

where $\boldsymbol{U}(t)$ is called the fundamental matrix solution, which is given by

$$\boldsymbol{U}(t) = \begin{bmatrix} 1 & 0 \\ \beta \sum\limits_{i=1}^{\infty} \frac{i}{k^2+(i\omega)^2}\left[\sqrt{\left(A_i^2+B_i^2\right)\left(k^2+(i\omega)^2\right)}\cos\Phi_i(t) - \left(i\omega A_i + kB_i\right)e^{-kt}\right] & e^{-kt} \end{bmatrix}.$$

Here, we introduce a constant matrix $\boldsymbol{B}$ and periodic matrix function $\boldsymbol{P}(t)$, which are defined as

$$\exp(\tau \boldsymbol{B}) \equiv \boldsymbol{U}(0)^{-1}\boldsymbol{U}(\tau),$$
$$\boldsymbol{P}(t) \equiv \boldsymbol{U}(t)e^{-t\boldsymbol{B}}.$$

$\boldsymbol{B}$ and $\boldsymbol{P}(t)$ for the model Eq (S1) are:

$$\boldsymbol{B} = \begin{bmatrix} 0 & 0 \\ \beta k \sum\limits_{i=1}^{\infty} \frac{i}{k^2+(i\omega)^2}\left(i\omega A_i + kB_i\right) & -k \end{bmatrix},$$

$$\boldsymbol{P}(t) = \begin{bmatrix} 1 & 0 \\ \beta \sum_{i=1}^{\infty} \frac{i}{k^2+(i\omega)^2} \left[ \sqrt{(A_i^2+B_i^2)(k^2+(i\omega)^2)} \cos \Phi_i(t) - (i\omega A_i + kB_i) \right] & 1 \end{bmatrix}. \tag{S5}$$

We denote the right and left eigenvectors of $\boldsymbol{B}$ as $\boldsymbol{\phi}$ and ${}^t\boldsymbol{\psi}$, respectively, where the superscript $t$ indicates transposition. According to the Floquet theory [2], the eigenvalues of $\boldsymbol{B}$ are called Floquet exponents. The eigenvalues of $\boldsymbol{B}$ for the model Eq (S1) are $\lambda_0 = 0$ and $\lambda_1 = -k$. That one of the exponents is zero is consistent with the fact that the considered model has a limit cycle. When $\lambda_0 = 0$, the eigenvectors are:

$$\boldsymbol{\phi}_0 = \begin{bmatrix} \omega \\ \beta \sum_{i=1}^{\infty} \frac{i\omega}{k^2+(i\omega)^2} (i\omega A_i + kB_i) \end{bmatrix}, \quad \boldsymbol{\psi}_0 = \begin{bmatrix} \frac{1}{\omega} \\ 0 \end{bmatrix}. \tag{S6}$$

When $\lambda_1 = -k$,

$$\boldsymbol{\phi}_1 = \begin{bmatrix} 0 \\ 1 \end{bmatrix}, \quad \boldsymbol{\psi}_1 = \begin{bmatrix} -\beta \sum_{i=1}^{\infty} \frac{i}{k^2+(i\omega)^2} (i\omega A_i + kB_i) \\ 1 \end{bmatrix}. \tag{S7}$$

According to the theory [1], the coefficient of variation (CV) of the oscillation period can be calculated using three components:

$$\mathrm{CV} = \frac{\epsilon}{\tau} \sqrt{R_{\Theta\Theta} + R_{\boldsymbol{hh}} + 2R_{\Theta\boldsymbol{h}}} + \mathcal{O}(\epsilon^2), \tag{S8}$$

where $R_{\Theta\Theta}$, $R_{\boldsymbol{hh}}$, and $R_{\Theta\boldsymbol{h}}$ represent the variance of the system-level phase, the autocorrelation of the amplitude deviation, and the cross-correlation between the system-level phase shift and the amplitude deviation, respectively. These three components can be expressed using $\boldsymbol{P}(t)$, $\boldsymbol{\phi}_i$, and $\boldsymbol{\psi}_i$ $(i = 1, ..., N-1)$ as follows:

$$\begin{aligned}
R_{\Theta\Theta} &= \int_0^{\tau} {}^t\boldsymbol{\psi}_0 \boldsymbol{P}(t)^{-1} \boldsymbol{G}[\boldsymbol{p}(t)]^2 \, {}^t\boldsymbol{P}(t)^{-1} \boldsymbol{\psi}_0 dt, \\
R_{\boldsymbol{hh}} &= \sum_{j=1}^{N-1} \sum_{k=1}^{N-1} [2 - \exp(\lambda_j \tau) - \exp(\lambda_k \tau)] \frac{[\boldsymbol{P}(t_{\mathrm{cp}})\boldsymbol{\phi}_j]_l [\boldsymbol{P}(t_{\mathrm{cp}})\boldsymbol{\phi}_k]_l}{[\dot{\boldsymbol{p}}(t_{\mathrm{cp}})]_l^2} \exp[(\lambda_j + \lambda_k) t_{\mathrm{cp}}] \\
&\times \left\{ \int_0^{t_{\mathrm{cp}}} \exp[-(\lambda_j + \lambda_k)t] \, {}^t\boldsymbol{\psi}_j \boldsymbol{P}(t)^{-1} \boldsymbol{G}[\boldsymbol{p}(t)]^2 \, {}^t\boldsymbol{P}(t)^{-1} \boldsymbol{\psi}_k dt \right. \\
&+ \left. \frac{\exp[(\lambda_j + \lambda_k)\tau]}{1 - \exp[(\lambda_j + \lambda_k)\tau]} \int_0^{\tau} \exp[-(\lambda_j + \lambda_k)t] \, {}^t\boldsymbol{\psi}_j \boldsymbol{P}(t)^{-1} \boldsymbol{G}[\boldsymbol{p}(t)]^2 \, {}^t\boldsymbol{P}(t)^{-1} \boldsymbol{\psi}_k dt \right\}, \\
R_{\Theta\boldsymbol{h}} &= \sum_{j=1}^{N-1} \exp(\lambda_j \tau) \frac{[\boldsymbol{P}(t_{\mathrm{cp}})\boldsymbol{\phi}_j]_l}{[\dot{\boldsymbol{p}}(t_{\mathrm{cp}})]_l} \\
&\times \int_0^{\tau} \exp(-\lambda_j t) \, {}^t\boldsymbol{\psi}_0 \boldsymbol{P}(t_{\mathrm{cp}} + t)^{-1} \boldsymbol{G}[\boldsymbol{p}(t + t_{\mathrm{cp}})]^2 \, {}^t\boldsymbol{P}(t_{\mathrm{cp}} + t)^{-1} \boldsymbol{\psi}_j dt,
\end{aligned} \tag{S9}$$

where $[\boldsymbol{x}]_l$ denotes the $l$-th element of vector $\boldsymbol{x}$. For $l = 1$ and 2, CV represents the fluctuation in the period of $\theta$ and $x$, respectively. By substituting Eqs (S2), (S5), (S6), and (S7) into Eq (S9) with $l = 2$, we obtain Eq (5).

### 2 Regulation by the different components in the Goodwin model Eq (1)

We considered the Goodwin model in which $w$ controls the downstream output in Fig 1 of the main text. We further examined alternative scenarios in which $u$ or $v$ control the downstream output (Fig S1). CV of the period depended on the choice of the variables and the value of $k$, and not on the values of $\alpha$ or $\beta$.

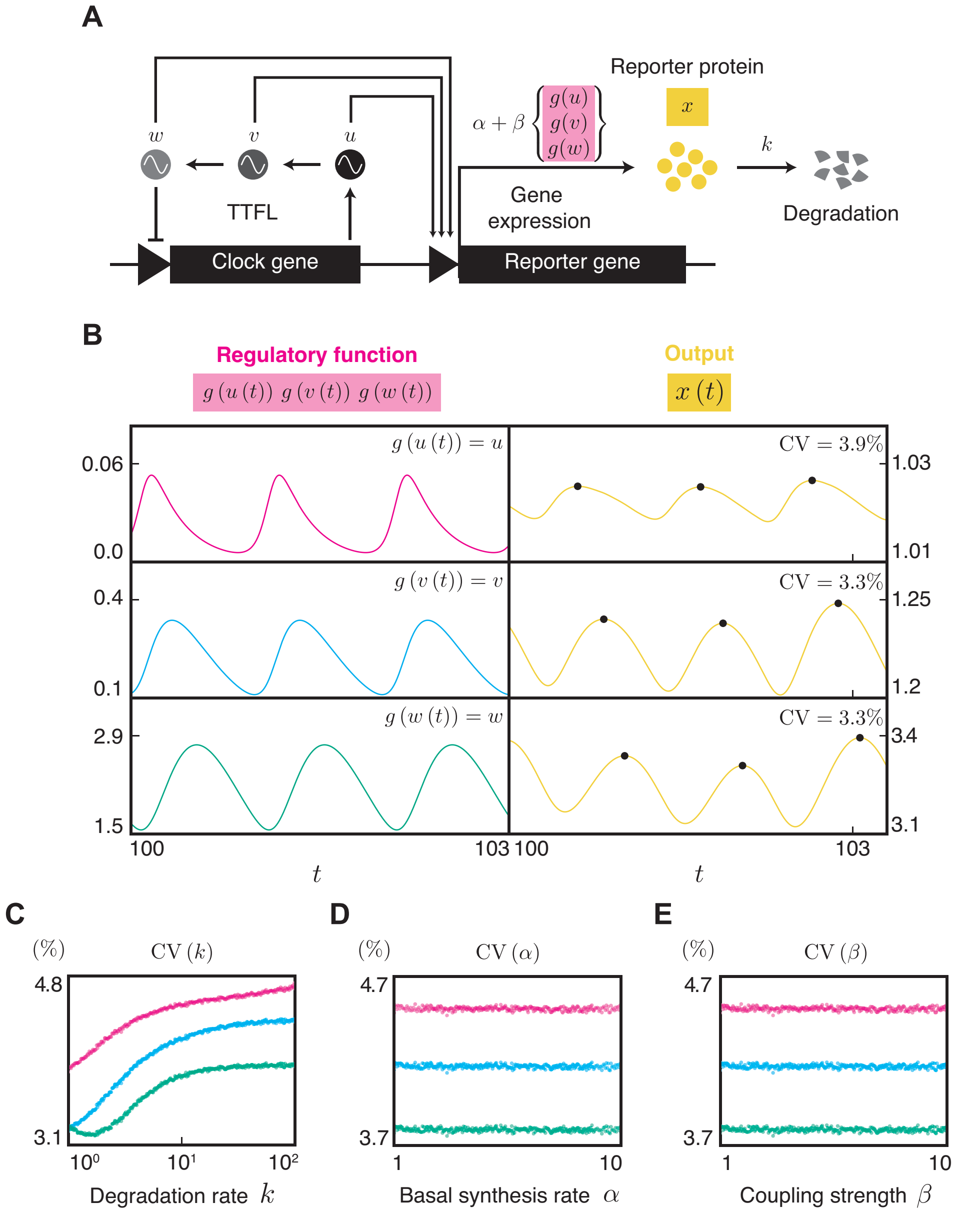


**Fig S1. Fluctuation of the output system driven by different variables in TTFL. (A)** Goodwin model representing TTFL coupled with a reporter system. Here, we assumed that one of the components $u$, $v$, and $w$ drives the reporter gene $x$. Gene expression noise induces fluctuations in $u$, as described in the main text. **(B)** The fluctuation of the output system depends on the choice of variable that regulates the output. The left column shows $u(t)$, $v(t)$, and $w(t)$ in the absence of noise. The right column shows $x(t)$ regulated by different variables $u$, $v$, or $w$. Black dots indicate the peaks of each cycle. $k_u = k_v = k_w = 0.1$, $\alpha = \beta = 1.0$, $k = 1.0$, $\epsilon = 10^{-4}$, $D = 1.0$, and $\tau = 39.7$. **(C)** The dependency of the degradation rate $k$ on the CV of the oscillation period. $\alpha = \beta = 1$. **(D)** The dependency of the basal synthesis rate $\alpha$ on the CV of the oscillation period. $\beta = 1$, $k = 10$. **(E)** The dependency of the coupling strength $\beta$ on the CV of the oscillation period. $\alpha = 1$, $k = 10$. The colors correspond to the functions in Fig B. The values of the other parameters are the same as in Fig B.

#### 3 Examples of regulatory functions obtained by Gibbs sampling

Fig S2 shows examples of waveforms of regulatory functions obtained by Gibbs sampling in Fig 5. The samples were randomly selected from each bin of the histogram.

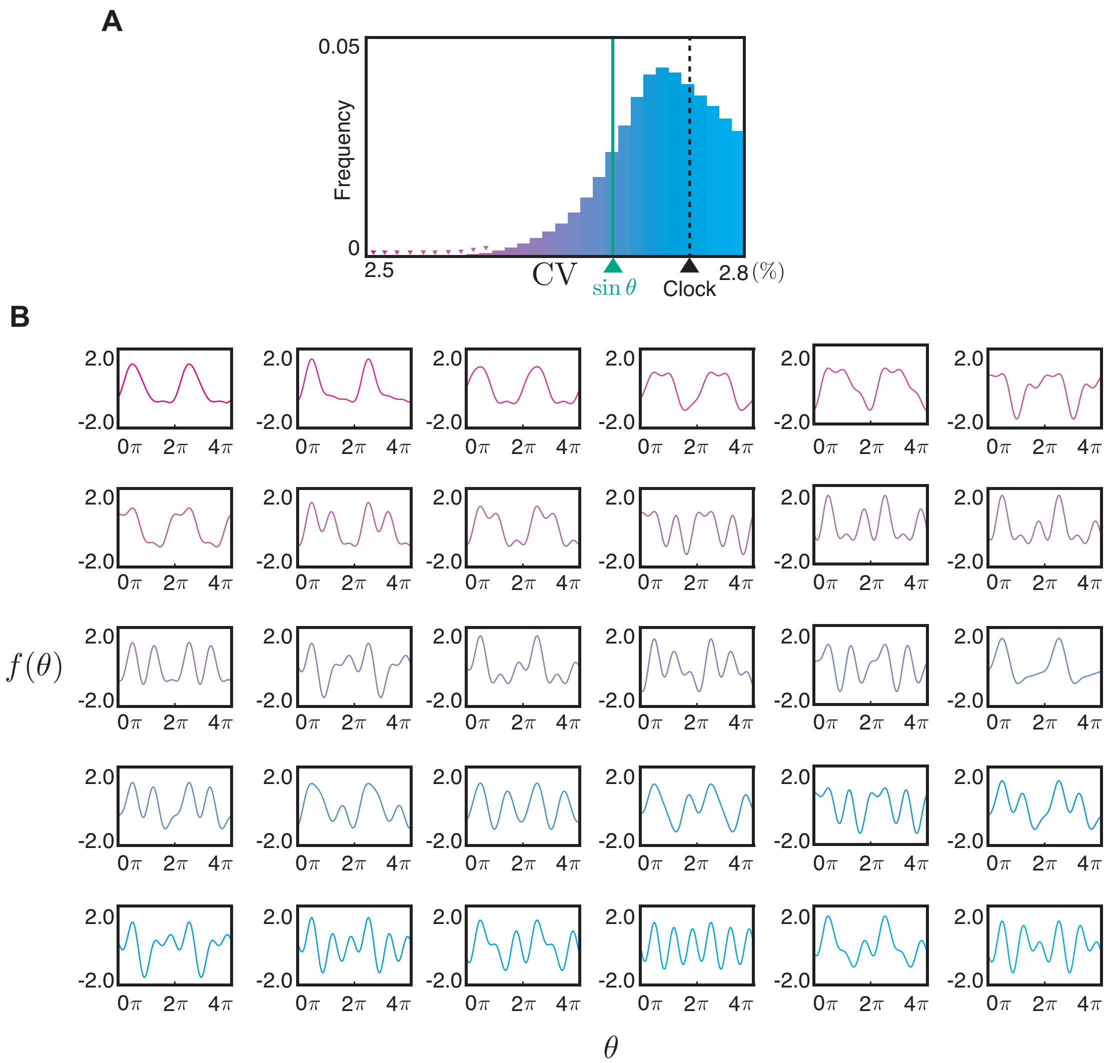


**Fig S2. Waveforms of the regulatory functions obtained by Gibbs sampling.** (**A**) Histogram of samples obtained from Gibbs sampling using Eq (5). The CV for the sinusoidal regulation, represented by the green line, is 2.7% (see §4 in the Supplementary text). The dashed line represents the CV of the clock, expressed as $\epsilon\sqrt{\frac{D}{2\pi\omega}}$ [3]. (**B**) Examples of waveforms selected from each bin of Fig A. Each curve corresponds to the bins of the histogram with the same color. It means that the CV of the waveforms increases from left to right within each row and from top to bottom across rows, with the smallest CV at the top-left and the largest CV at the bottom-right.

### 4 Methods to obtain CV of period

We used two methods to obtain CV in this paper: one is using the numerical data of the oscillation period measured in simulations. The other method calculates CV based on Eq (5), which requires the values of the parameters in the model described by Eq (2) and the time of the checkpoint $t_{\mathrm{cp}}$.

#### 4.1 CV obtained from numerical simulations

We measured the peak-to-peak intervals of the time series obtained by Euler-Maruyama method with the step size $\Delta t = 0.001$ for Figs 1-3, S1, and S5. For Fig 4, the intervals were measured based on checkpoints. Suppose that we numerically observed a time series of a variable $\mathsf{x}$, which is denoted as $\hat{\mathsf{x}}(i\Delta t)$, where $i = 0, 1, 2, \ldots$ We collected 1000 intervals from the time series of $\hat{\mathsf{x}}(i\Delta t)$ to calculate a CV value. The mean CV was computed as the average of 100 repetitions of CV measurement. Any interval outside the range of $[0.8\tau, 1.2\tau]$, where $\tau$ is the period of a noiseless system, was excluded from the calculation of CV as an outlier.

We detected the peak time in the $j$-th cycle, $t^j_{\mathrm{peak}}$ by searching for the local maximum of $\hat{\mathsf{x}}(i\Delta t)$ with the highest value within a cycle (Fig S3A). We checked local maximality of a point by looking at $\hat{\mathsf{x}}(i\Delta t)$ in the range of $[(i-10)\Delta t, (i+10)\Delta t]$. For $t^1_{\text{peak}}$, we discarded $0 \le i\Delta t < 100$ to avoid dependence of the initial condition and searched for local maxima within the range $100 \le i\Delta t \le 105$. The highest point in all collected local maxima within this range was defined as $t^1_{\text{peak}}$. For $j \ge 2$, $t^j_{\text{peak}}$ was searched within $[t^{j-1}_{\text{peak}} + 0.8\tau, t^{j-1}_{\text{peak}} + 1.2\tau]$. If no peaks were found, the search range was repeatedly expanded by $0.4\tau$ until the next peak was detected. These processes were repeated until the end of the simulation or until 1200 peaks were obtained.

On the other hand, we detected the checkpoint time in $j$-th cycle, $t^j_{\text{cp}}$, based on the combination of a threshold, $x_{\text{cp}}$, and the gradient of $\mathsf{x}$ when crossing $x_{\text{cp}}$; that is, we should take into account of the increase or decrease in $\hat{\mathsf{x}}(i\Delta t)$ (Fig S3B). For the numerical simulation shown in Fig 4, $x_{\text{cp}}$ was set to the midpoint of the oscillation range of $p_2(t)$, and the crossing direction was upward. In some cases, $\hat{\mathsf{x}}(i\Delta t)$ may cross $x_{\text{cp}}$ two or more in the same direction within a single cycle. To uniquely determine the checkpoints for each cycle, we detected all points that satisfy $\mathsf{x}(t) = x_{\text{cp}}$. These candidate points were divided into groups based on the troughs of $\hat{\mathsf{x}}(i\Delta t)$. The earliest candidate in the $j$-th group was defined as the time of the checkpoint $t^j_{\text{cp}}$. We first detected the time sequences of the troughs of $\hat{\mathsf{x}}(i\Delta t)$, $t^j_{\text{trough}}$, which were detected with the same procedure as peak detection. Then we searched for the checkpoint times, $t^j_{\text{cp}} \equiv \min_i(t^j_{\text{trough}} + i\Delta t)$ under the constraint of $\hat{\mathsf{x}}(t^j_{\text{trough}} + i\Delta t) \ge x_{\text{cp}}$.

$\hat{\mathsf{x}}(i\Delta t)$ ···

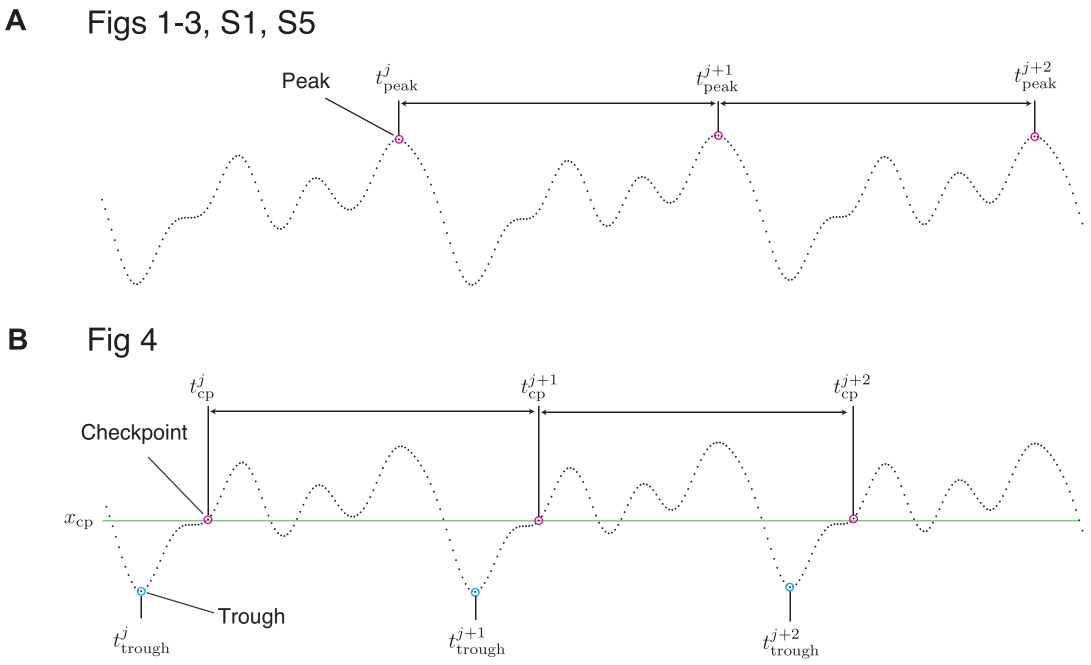


**Fig S3. Methods of measuring periods through simulation.** (**A**) Measuring CV of periods based on peaks. We detected the peaks of each cycle. The time interval between adjacent peaks is defined as a period. (**B**) Methods of measuring periods based on checkpoints.

#### 4.2 CV obtained from Eq (5)

Eq (4) in the main text suggests that the CV of the oscillation period is calculable based on three components described by Eq (5). The assignment of all the values of parameters and $t_{\text{cp}}$ in Eq (5) yields a CV value.

We determined the value of $t_{\text{cp}}$ in several ways. For Fig 4, we first calculated the value of threshold $x_{\text{cp}}$ as the midpoint of the oscillatory range, i.e., $x_{\text{cp}} = (\min_t p_2(t) + \max_t p_2(t))/2$ (Fig S4A). In addition, we obtained the value of $t_{\text{trouph}} \equiv \min_{0 \le t < 1} p_2(t)$. Then, we numerically found the value of $t_{\rm cp} \equiv \min_{i\in\mathbb{N}}(t_{\rm trough}+0.001i)$ under the constraints that $p_2(t_{\rm trough}+i\varepsilon) \geq x_{\rm cp}$ and $\frac{\dot{p}_2(t_{\rm cp})}{|\dot{p}_2(t_{\rm cp})|} = 1$.

For DE in Fig S6, The values of $x_{\rm cp}$ were points equally spaced between the maximum and minimum values of the limit cycle (See §5 and Fig S4B). Then, we numerically found the value of $t_{\rm cp} \equiv \min_{i\in\mathbb{N}}(0.001i)$ under the constraints that $p_2(i\varepsilon) \geq x_{\rm cp}$ and $\frac{\dot{p}_2(t_{\rm cp})}{|\dot{p}_2(t_{\rm cp})|} = 1$.

For the Gibbs sampling based on the analytical expression (Fig 5), we set $t_{\rm cp} = 0$ (Fig S4C). For $\sin\theta$, we set $t_{\rm cp} = 0, 0.001, 0.002, ..., 0.999$ and then chose a median of 1000 obtained CV.

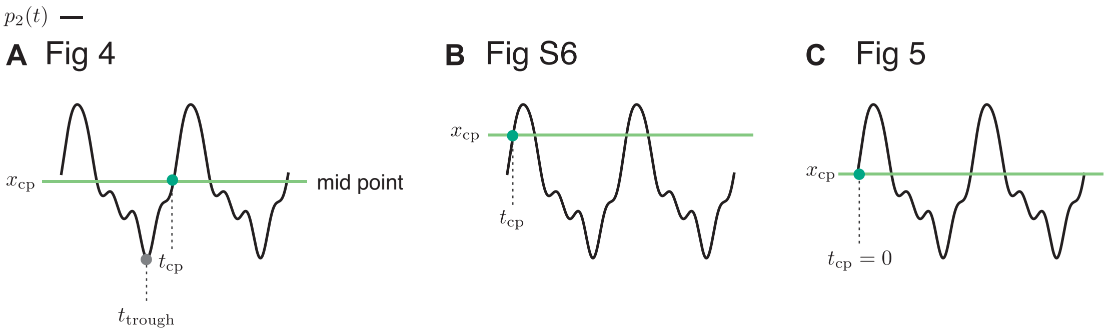


**Fig S4. The checkpoint $t_{\rm cp}$ for a threshold $x_{\rm cp}$.** (**A**) Detecting $t_{\rm cp}$ for the midpoint as a threshold. (**B**) Detecting $t_{\rm cp}$ for a given threshold. (**C**) The checkpoint time was set to $t_{\rm cp} = 0$.

### 5 Optimization of the regulatory function via an evolutionary algorithm

We employed an evolutionary algorithm in a simulation to determine the regulatory function $f$ that minimizes the CV of the oscillation period. We adopted Differential Evolution (DE) with a /rand/1/bin schema [4]. For the hyperparameters of DE, the population size was 50, the mutation factor was 0.9, and the crossover probability was 0.8.

We optimized two types of fitness functions. The first fitness function was the average of 100 CV values, where each CV was calculated from 1000 cycles measured by the peak-to-peak interval of numerical simulation of Eq (2) under a fixed value of $k$. The DE identified an optimal rhythmic regulatory function with a CV smaller than $\sin\theta$. The optimized function closely resembles a sine curve, supporting the idea that sine-like functions can effectively reduce fluctuations (Fig S5).

Next, we optimized the averaged CV, denoted as $\langle \mathrm{CV} \rangle_{\mathrm{cp},k} \equiv \frac{1}{N_k N_{\rm cp}} \sum_{i=0}^{N_k-1} \sum_{j=0}^{N_{\rm cp}-1} \mathrm{CV}\left(f, k_i, x_{\rm cp}^j\right)$, where $N_{\rm cp}$ is the number of different thresholds $x_{\rm cp}$, and $N_k$ is the number of different values of $k$. $x_{\rm cp}^j$ are the threshold points equally spaced between the maximum and minimum values of the limit cycle, defined as $\min_t p_2(t) + \left[\max_t p_2(t) - \min_t p_2(t)\right] \cdot \left(0.2 + \frac{0.6j}{N_{\rm cp}-1}\right)$. The values of $k_i$ are equally spaced on a logarithmic scale between $10^0$ and $10^2$, i.e., $k_i = 10^{\frac{2i}{N_k-1}}$. We set $N_k = N_{\rm cp} = 10$. The DE iteration procedure halted when the number of generations reached $2\times10^5$ or when the change in CV between generations was less than $10^{-7}$ for $4 \times 10^4$ iterations. The DE algorithm again identified the more robustly denoising regulatory function compared with $\sin\theta$. The fluctuation of the optimal function was significantly smaller than that of the sinusoidal function in the range $10^0 \leq k \leq 10^1$. In contrast to the former fitness function, the optimized function did not resemble a sinusoidal function (Fig S6).

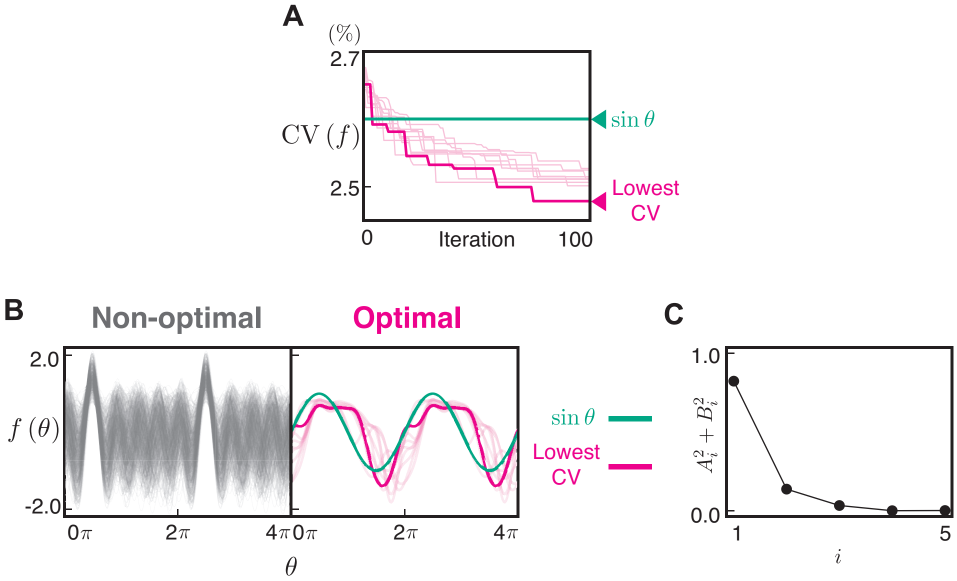


**Fig S5. Optimization of regulatory function for CV.** (**A**) The convergence curves of CV in the optimization with Differential Evolution. Ten trials are displayed, with the thick line representing the trial with the lowest CV at the 100th iteration. The parameters for the numerical simulation were $\omega = 2\pi$, $\epsilon = 0.1$, $D = 3$, $\alpha = 3$, $\beta = 1$, $k = 10$, and $f = \sum_{i=1}^{5} (A_i \cos(i\theta) + B_i \sin(i\theta))$. The values of Fourier coefficients for the function giving the lowest CV are displayed in Table S1. The CV for the sinusoidal regulation $f(\theta) = \sin\theta$, represented by the green line, is 2.6%. (**B**) The waveform of randomly selected and optimized regulatory functions $f$. The left panel shows the initial population of the optimization of $f$, and the right panel shows 10 examples of the optimized $f$. All waveforms are aligned to have their maximum value at $\theta = \frac{\pi}{2}$. (**C**) Power of the Fourier coefficients for the function exhibiting the lowest CV.

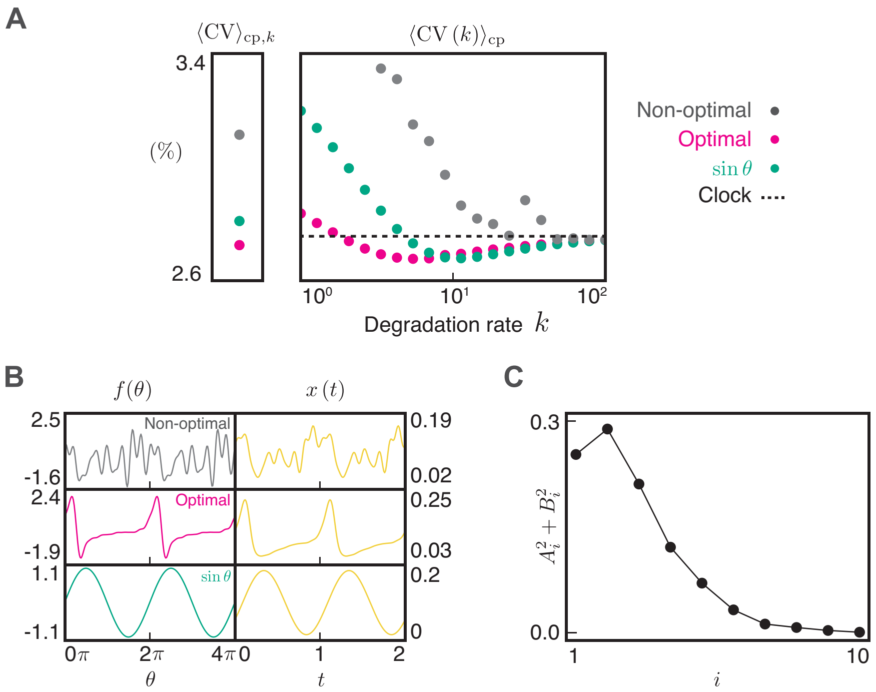


**Fig S6. Robustly denoisable regulatory function.** (**A**) The DE algorithm identified the regulatory function that can robustly reduce avraged fluctuations across a wide range of $x_{\mathrm{cp}}$ and $k$. The values of $\langle \mathrm{CV} \rangle_{\mathrm{cp},k}$ and

$$\langle \mathrm{CV}\,(k) \rangle_{\mathrm{cp}} \equiv \frac{1}{N_{\mathrm{cp}}} \sum_{i=1}^{N_{\mathrm{cp}}} \mathrm{CV}\left(f, k, x_{\mathrm{cp}}^{i}\right)$$

for the non-optimal function, the optimal function, and $\sin\theta$ are displayed, respectively. The values of Fourier coefficients for the optimal and non-optimal functions are displayed in Table S1. The parameters for the numerical simulation of Eq (2) were $\omega = 2\pi$, $\epsilon = 10^{-1}$, $D = 3$, $\alpha = 4$, and $\beta = 1$, $N_k = N_{\mathrm{cp}} = 20$. The regulatory function were represented as

$$f = \sum_{i=1}^{10} \left(A_i \cos\left(i\theta\right) + B_i \sin\left(i\theta\right)\right).$$

(**B**) Waveforms of the regulatory function $f(\theta)$. The dynamics of the output system $x(t)$ produced by each regulatory function are displayed on the right side of the corresponding $f$. (**C**) Power of the Fourier coefficients for the optimal function.

### 6 Supplementary table

**Table S1. The values of Fourier coefficients for the function $f(\theta)$ in the figures.**

| Figure | $n$ | $A_n$ | $B_n$ |
|---|---|---|---|
| Fig 2B red | 1 | −0.1281 | 0.4668 |
| | 2 | 0.1392 | −0.498 |
| | 3 | −0.4538 | 0.4919 |
| | 4 | 0.219 | −0.0376 |
| | 5 | 0.0127 | 0.0292 |
| Fig 2B blue | 1 | 0.5329 | −0.2926 |
| | 2 | 0.0824 | −0.1728 |
| | 3 | −0.4913 | −0.029 |
| | 4 | 0.5426 | 0.1671 |
| | 5 | −0.1551 | −0.0714 |
| Fig 4 red | 1 | 0.2074 | −0.87 |
| | 2 | −0.0591 | −0.0953 |
| | 3 | −0.0372 | −0.0862 |
| | 4 | −0.0841 | −0.0632 |
| | 5 | 0.073 | 0.1345 |
| | 6 | −0.061 | −0.1741 |
| | 7 | 0.0657 | −0.128 |
| | 8 | −0.1578 | 0.0889 |
| | 9 | −0.1763 | 0.0208 |
| | 10 | −0.0645 | 0.1446 |
| Fig 4 blue | 1 | −0.2354 | 0.0473 |
| | 2 | −0.3604 | 0.1296 |
| | 3 | 0.0763 | −0.2161 |
| | 4 | 0.3716 | −0.0277 |
| | 5 | −0.0897 | −0.1321 |
| | 6 | 0.0039 | 0.3843 |
| | 7 | −0.1749 | −0.1616 |
| | 8 | −0.098 | −0.0056 |
| | 9 | 0.2348 | −0.1892 |
| | 10 | 0.4144 | −0.3195 |

| Figure | $n$ | $A_n$ | $B_n$ |
|---|---|---|---|
| Fig S5B red | 1 | −0.8816 | 0.2163 |
| | 2 | −0.243 | 0.2813 |
| | 3 | 0.0 | 0.1848 |
| | 4 | −0.0347 | 0.0062 |
| | 5 | −0.0165 | −0.0459 |
| Fig S6B grey | 1 | 0.2084 | −0.2573 |
| | 2 | 0.1857 | −0.3164 |
| | 3 | 0.0278 | 0.1809 |
| | 4 | −0.1702 | 0.3381 |
| | 5 | −0.1826 | 0.2263 |
| | 6 | 0.2657 | −0.0222 |
| | 7 | −0.3183 | −0.0837 |
| | 8 | 0.0906 | 0.2103 |
| | 9 | 0.258 | 0.2142 |
| | 10 | 0.3587 | −0.1465 |
| Fig S6B red | 1 | 0.4673 | −0.1856 |
| | 2 | 0.5089 | 0.1731 |
| | 3 | 0.254 | 0.3826 |
| | 4 | −0.0709 | 0.3409 |
| | 5 | −0.2232 | 0.143 |
| | 6 | −0.1679 | −0.0645 |
| | 7 | −0.0286 | −0.1069 |
| | 8 | 0.0499 | −0.0696 |
| | 9 | 0.0547 | −0.0122 |
| | 10 | 0.0219 | 0.0131 |
